# Synaptotoxic effects of extracellular tau are mediated by its microtubule-binding region

**DOI:** 10.1101/2025.02.21.639417

**Authors:** Tomas Ondrejcak, Neng-Wei Hu, Emily Coode, Tom Campbell, Grant T. Corbett, Ivan Doykov, Kevin Mills, Dominic M. Walsh, Frederick J. Livesey, Michael J. Rowan, Igor Klyubin

## Abstract

Immunotherapies targeting extracellular tau share the premise that interrupting cell-to-cell spread of tau pathology in Alzheimer’s disease (AD) will slow dementia pathogenesis. How these interventions affect the actions of synaptotoxic, extracellular tau species that may help mediate cognitive impairment is relatively unknown. Here, we assayed synaptic plasticity disruption in anaesthetised live rats caused by intracerebral injection of synaptotoxic tau present either in (a) secretomes of induced pluripotent stem cell-derived neurons (iNs) from people with Trisomy 21, the most common genetic cause of AD, or (b) aqueous extracts of human AD brain. Extracellular tau in iN secretomes was found to include fragments that contain the extended microtubule binding regions of tau, MTBR/R’ and adjacent C-terminal peptides. Immunodepletion or co-injection with antibodies targeting epitopes within these fragments prevented the acute disruption of synaptic plasticity by these patient-derived synaptotoxic tau preparations. Conversely, a recombinant human tau fragment encompassing the core MTBR/R’- region present in tau fibrils, tau297-391 potently mimicked this deleterious action of patient-derived tau. MTBR/R’-directed antibodies also rapidly reversed a very persistent synaptotoxic effect of soluble brain tau. Our findings reveal a hitherto relatively unexplored potential benefit of targeting MTBR/R’.

## Introduction

Although great strides have been made in understanding the physiology and pathology of the microtubule-associated protein, tau, we still lack successful tau-based therapeutic interventions. Some of the most advanced tau-targeting trial therapies for Alzheimer’s disease (AD) are based on use of antibodies intended to bind and neutralize extracellular tau species capable of propagating tau pathology. Besides the potential spreading of tau aggregation between neurons, there is evidence that certain forms of extracellular tau have additional pathogenic properties, including disrupting synaptic function [12, 13].

In human brain there are six different splice isoforms of tau that give rise to proteins of 352-441 amino acids, all of which undergo posttranslational modifications including truncation. C-terminal (CT) fragments are mainly found inside, whereas N-terminal (NT) and mid-region (MR) fragments predominate outside neurons. Only fragments which contain the microtubule-binding region (MTBR) are competent to aggregate [5]. Clinical trials targeting the NT of tau have been unsuccessful and focus has shifted to testing antibodies which target epitopes in or near the MTBR. Tau fragments that include the MTBR and the CT adjacent pseudorepeat R’(aa369-399) [8] (referred to as the MTBR/R’ domain of tau), are enriched in tauopathy brain including AD [1, 26]. Indeed, the accumulation of certain MTBR fragments in the brain is correlated with cognitive impairment [1, 11, 26] providing support for investigating MTBR targeting strategies. More detailed understanding of the role of soluble extracellular MTBR-containing tau fragments in AD should inform planned clinical trials.

AD is characterised by synaptic failure. To date, little is known about the vulnerability of synapses to pathologically relevant forms of MTBR-containing tau fragments. Several preparations are available to investigate the synaptotoxicity of tau including recombinant tau, patient-derived brain samples and secretomes from induced pluripotent stem cell (iPSC)-derived neurons (iN). Previously, we reported that removing tau from aqueous brain extracts of some AD patients, prevented inhibition of long-term potentiation (LTP), an electrophysiological correlate of memory [17-19]. In addition, we discovered that the secretomes of iNs from people with Down syndrome (Trisomy 21, Ts21), the most common cause of early onset AD [15], also disrupt synaptic plasticity in a tau-dependent manner [12]. Both patient-derived preparations contain a range of tau species. Secretomes from iNs, like human CSF, have abundant N-terminal and mid-region fragments of tau with relatively low levels of MTBR-containing tau [3, 7, 12, 23], whereas aqueous extracts of AD brain contain mostly intracellular full length and MTBR-containing tau.

Here, we studied the involvement of extended MTBR/R’-containing tau species [5] in the acute inhibition of LTP by patient-derived soluble synaptotoxic tau in anaesthetized rats *in vivo*. We also tested the ability of two anti-tau antibodies targeting MTBR/R’ and the adjacent CT region, Gen2B and Gen2A, respectively, to reverse a persistent disruption of synaptic plasticity induced by AD brain tau.

## Materials and methods

See Supplementary materials for full details.

### Generation of cortical cultures and secretome collection

The iPSC lines used in this study were non-demented control (NDC) [14], and Ts21 [20]. Ts21 iPSCs were generated from two individuals with Down syndrome (referred to as Line1 and Line2) using the CytoTune-iPS 2.0 Sendai Reprogramming Kit (ThermoFisher). Human cortical neurons were produced from all iPSC lines essentially as described [24].

### Human brain tissue

Frozen tissue was obtained from two cases of end-stage AD (referred to as AD1 and AD6) [17-19].

### Custom MSD (Meso Scale Diagnostics) assay

To quantify extracellular levels of MTBR/R’-containing tau fragments, a custom MSD electrochemiluminescence assay was developed using uncoated MULTI-ARRAY 96-well SECTOR plates (MSD), Gen2B (epitope in 369-381) and Tau5 (epitope in 218-225) antibodies (Fig.ure 1A).

**Fig. 1.**
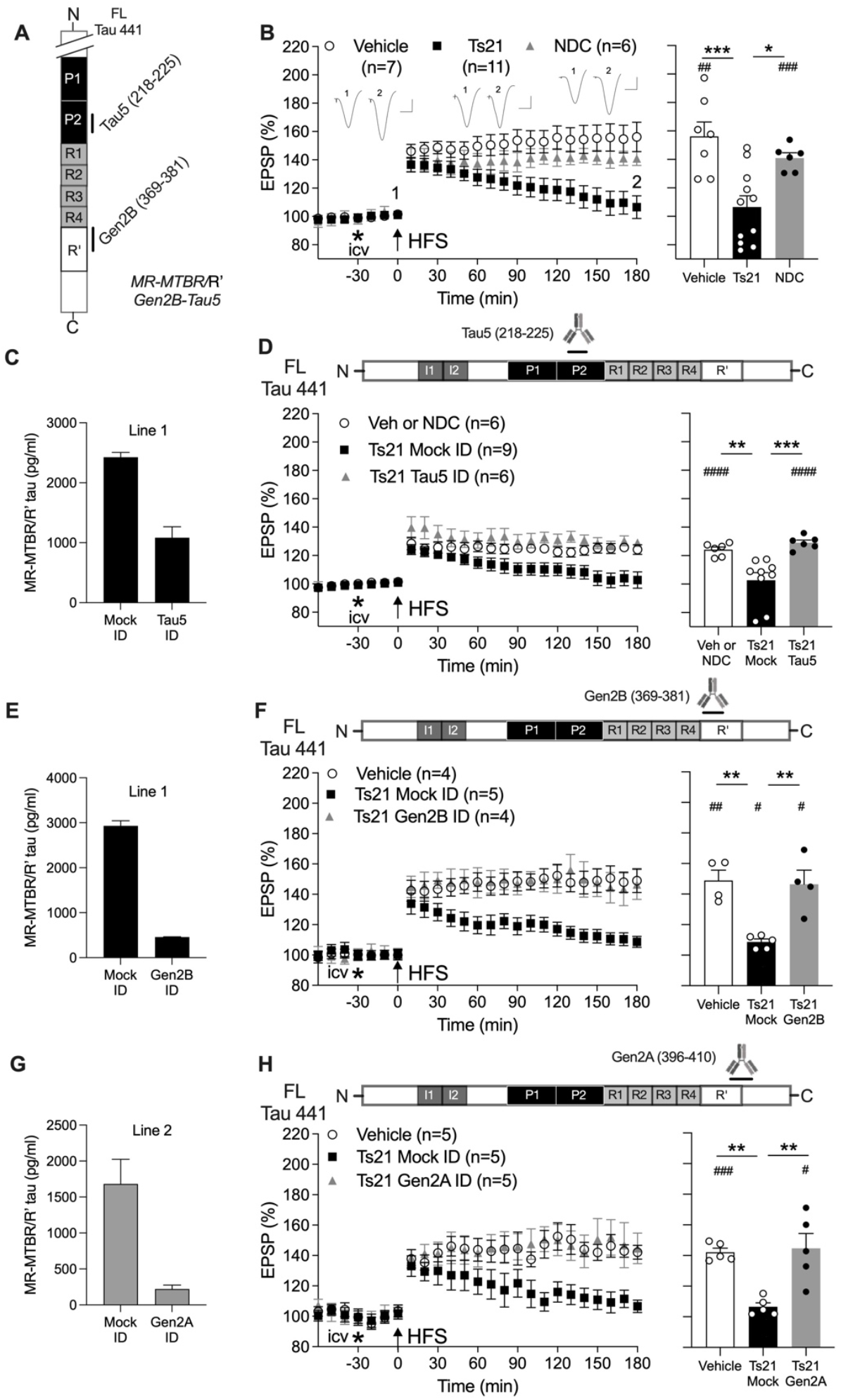
MTBR/R’-containing extracellular tau fragments secreted by Ts21 iNs are synaptotoxic. (**A**) Antibodies used to capture/detect MR-MTBR/R’ tau fragments in a custom Meso Scale Diagnostics assay (**C, E**, and **G**). (**B**) High-frequency stimulation (HFS, arrow) triggered robust LTP of hippocampal synaptic transmission in anaesthetized (urethane, 1.6 g/kg, i.p.) rats after intracerebroventricular (i.c.v, asterisk) injection of either vehicle or the secretome from non-demented control (NDC) in one of two lines of iPSC-derived neurons (iNs). In contrast, LTP decayed to baseline 3 h after application of HFS in animals injected with Ts21 secretomes. (**C**) Immunodepletion (ID) with the monoclonal MR-directed antibody Tau5 lowered MTBR/R′ tau level by ∼55%. (**D**) After ID with Tau5 Ts21 secretomes no longer inhibited LTP. (**E-H**) ID either with the monoclonal antibody Gen2B, directed at R’, (**E** and **F**), or Gen2A, directed at R’ and the adjacent CT, (**G** and **H**), caused a >85% reduction of MR-MTBR/R’ fragment concentration, and prevented the inhibition of LTP by Ts21 secretomes. In **B, D, F** and **H**, left-hand panels show the time course of LTP. Summary bar charts of LTP magnitude during the last 10 min are displayed in the right-hand panels. Insets in **A** show representative field EPSP traces at the times indicated. Calibration bars: vertical, 1 mV; horizontal, 10 ms. In **B, D, F** and **H**, values are mean ± SEM. ^#^p < 0.05, ^##^p < 0.01, ^###^p < 0.001, ^####^p < 0.0001 compared with pre-HFS, paired *t*-test; *p < 0.05, **p < 0.01, ***p < 0.001, one-way ANOVA followed by Bonferroni’s multiple-comparison tests.

### Animals

Most of experiments were conducted on adult male Lister Hooded rats as described previously [18]. Intracerebroventricular (i.c.v.) injections were carried out under either recovery (ketamine and medetomidine, 60 and 0.4 mg/kg, respectively, i.p.) or non-recovery (urethane, 1.6 g/kg, i.p.) anaesthesia.

### Mass spectrometry (MS)

Liquid chromatography with tandem MS analysis, similar to that described previously [4], was used to investigate MTBR-containing tau species in size exclusion chromatography (SEC) fractions of iN secretomes..

### Data analysis

Electrophysiological data are expressed as the average EPSP amplitude during the last 10 min epoch before and 170–180 min after HFS. Sample sizes were chosen based on our previous publications [17, 18]. The ability to induce LTP within each group was assessed *a priori* using paired two-tailed *t* tests. Differences in the magnitude of potentiation between groups were analyzed using one- or two-way ANOVA with Bonferroni’s *post hoc* tests or by unpaired two-tailed *t* tests, as appropriate. A *p* value of <0.05 was considered statistically significant. Statistical analyses were performed in GraphPad Prism software (10.4.1). For immunoassays, bars represent mean of two technical replicates ± SD.

## Results

### MTBR/R’-containing tau fragments in Ts21 iN secretomes mediate inhibition of hippocampal LTP

First, we explored the role of MTBR/R’-containing fragments in the synaptic plasticity-disrupting action of extracellular tau secreted by Ts21 iNs. The most abundant tau fragments in Ts21 secretomes are recognized by an antibody targeting the MR of tau (aa218-225), Tau5 [12]. Consistent with our previous report [12] i.c.v. injection of Ts21, but not NDC, iN secretomes robustly inhibited hippocampal LTP in anaesthetised rats (Fig. 1B). Interestingly, using different tau immunoassays (see Fig. 1A and Fig. 1S A), we found that Tau5 ID not only reduced the MR but also, to a lesser extent, the MR-MTBR/R’ and CT pools of Ts21 tau fragments (Fig. 1C, Fig. 1S A). Whereas LTP inhibition by Ts21 iN secretomes was prevented by ID with Tau5 (Fig. 1D), partial ID with an antibody directed to the extreme CT of tau (aa404-441), Tau46, appeared ineffective (Fig. 1S B). In contrast, ID with K9JA, a polyclonal antibody recognising a wide epitope including the MTBR and CT of tau (aa243-441), had similar effects to ID with Tau5. Thus, K9JA ID of Ts21 iN secretomes reduced CT tau by ∼85%, with a modest effect on MR tau (∼30%) when compared with Mock ID with pre-immune serum (PIS) and fully abrogated the LTP deficit (Fig. 1S C,D).

To investigate whether regions C-terminal to the MTBR were present in synaptotoxic forms of tau, we probed the region of tau spanning the R’ and adjacent CT sequence using two novel monoclonal antibodies, Gen2B and Gen2A. Both Gen2B, which is directed at the R’ domain (aa369-381) and Gen2A (targeting R’ and the adjacent CT region, aa396-410), were very effective in ID of MR-MTBR/R’ tau in Ts21 iN secretomes (>85% reduction in both cases, Fig. 1F,E). Importantly, the magnitude of LTP was indistinguishable from controls when animals were injected with either Gen2B ID (Fig. 1G) or Gen2A ID (Fig. 1H) secretomes. Complementing the ID approach, we co-injected some animals with Gen2A to neutralise tau fragments containing the CT sequence proximal to the MTBR. These animals maintained normal LTP after i.c.v administration of a mixture of Ts21 secretome and Gen2A mAb (Fig. 2S).

**Fig. 2.**
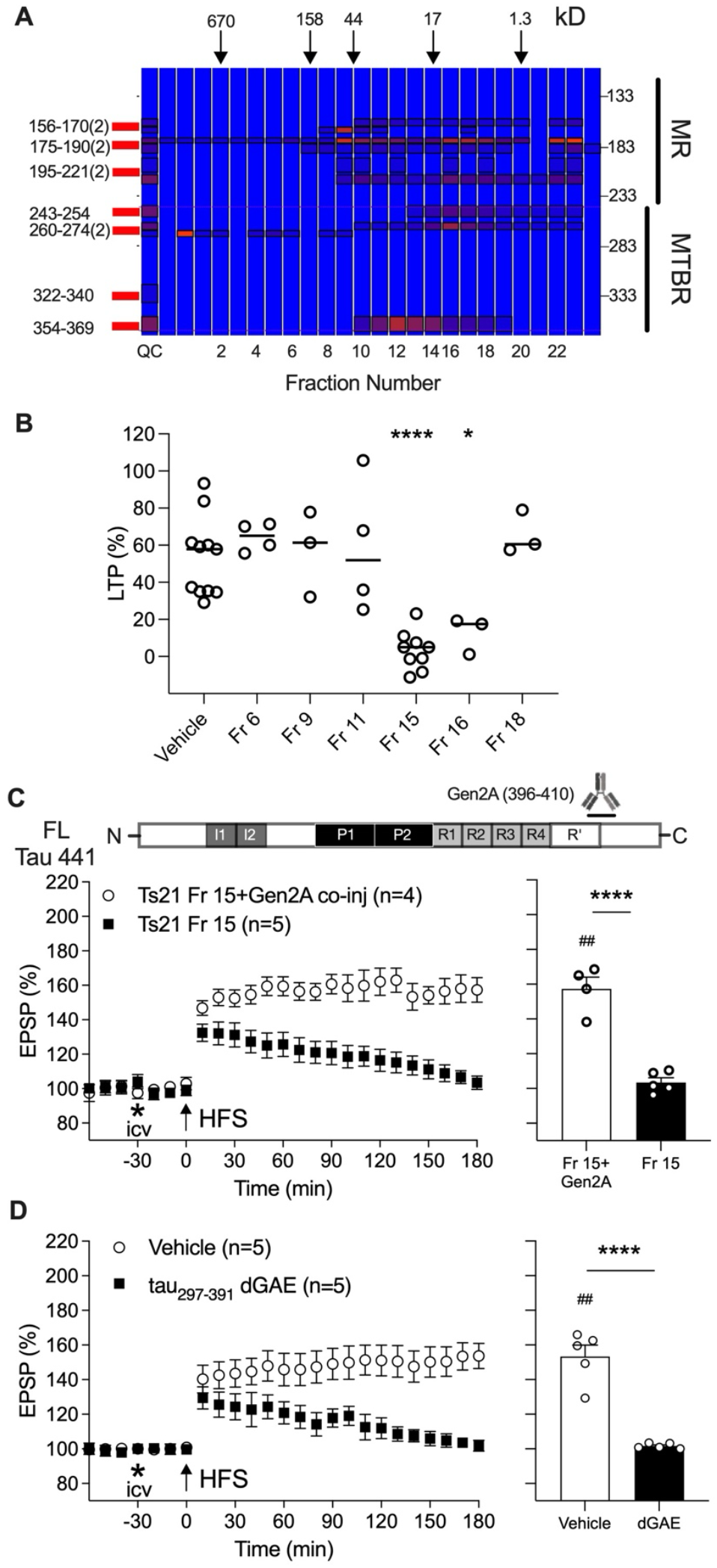
Tau fragments in certain Ts21 iN secretome SEC fractions potently inhibit LTP in an MTBR/R’-CT-dependent manner. (**A**) Mass spectrometry (MS) analysis of MR and MTBR tau in Ts21 iN secretome fractionated using Size Exclusion Chromatography (SEC). Tau fragments containing MTBR appear to be most abundant in fractions 12-14 and 16-17. Note that insufficient quantity of fraction 15 was available following the functional LTP experiments to analyse by MS. The elution of globular protein standards is indicated by arrows. Color intensity reflects peptide abundance. Quality control (QC) sample comprises all fractions analyzed (see Supplementary Fig. 5). Data from adjacent fragments containing unique tryptic peptides were combined (2). (**B**) In anaesthetized rats i.c.v. injection of Ts21 Mock ID SEC fractions 15 (Fr 15) and 16 (Fr 16) significantly inhibited hippocampal LTP. Circles represent individual animal values. *p < 0.05, ****p < 0.0001, compared with Vehicle group, one-way ANOVA followed by Bonferroni’s multiple-comparison tests. (**C**) Co-injection of Gen2A mAb (2.5 µg per injection) prevented the inhibition of LTP by Fr 15. (**D**) The recombinant human tau fragment tau297-391 (also known as dGAE fragment[16]) (80 pg/injection i.c.v.) potently inhibited LTP. This dose did not affect baseline synaptic transmission in the absence of HFS conditioning stimulation (pre-vs 210 min post-injection, n=3, p=0.3778, data not shown). Importantly, the amount of tau297-391 used was similar to MR-MTBR/R’-containing fragments in the Ts21 unfractionated secretomes (34-60 pg/injection i.c.v., Fig. **1C**,**E**,**G**). Left-hand panels in **C, D** show the time course of LTP. Summary bar charts of LTP magnitude during the last 10 min are displayed in the right-hand panels. Values are mean ± SEM. ^##^p < 0.01, compared with pre-HFS, paired *t*-test; ****p < 0.0001, unpaired *t*-test tests.

To further study the nature of Ts21 iN-derived synaptotoxic tau fragments, we used Ts21 secretomes that had been fractionated by SEC [12]. Fig. 3S outlines the distribution of MR and CT tau fragments across the fractions determined using immunoassays, consistent with what was described by us previously [12]. The relative abundance of MTBR was assessed using semi-quantitative MS and found especially in fractions 12-17 (Fig. 2A).

**Fig. 3.**
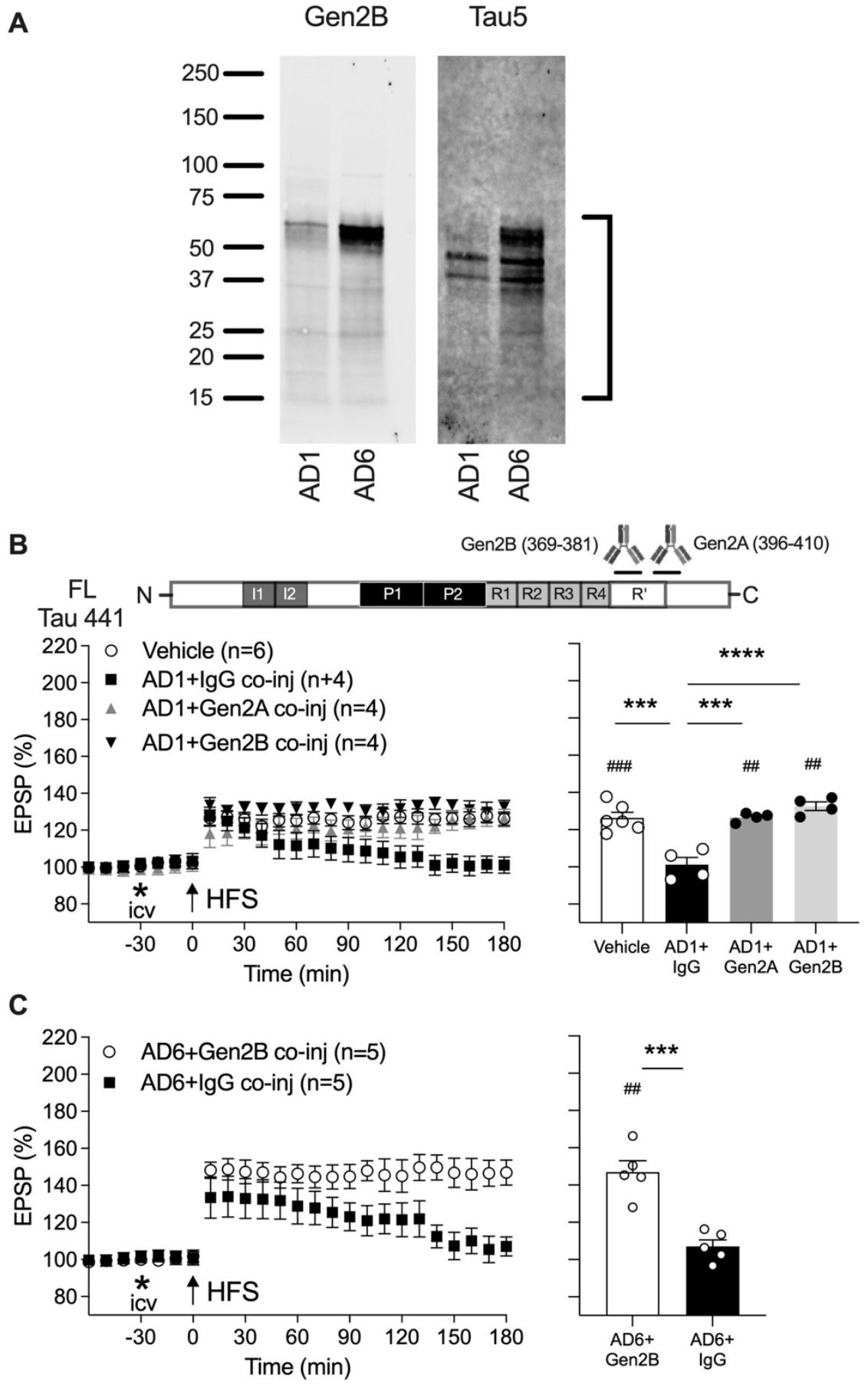
MTBR/R’ is required for acute plasticity disruption by synaptotoxic tau in AD brain extracts. (**A**) Aqueous extracts from the brains of two people with AD (AD1 and AD6) were analysed by Western blotting using the monoclonal anti-tau antibodies Gen2B and Tau5. Migration of SDS-PAGE molecular weight standards (in kDa) is indicated on the left and bracket on the right highlights the position of certain tau species. It should be noted that AD6 extract had much higher total tau concentration as measured by immunoassay[18]. (**B**) Unlike isotype control IgG, anaesthetized rats co-injected with AD1 extract and either Gen2A or Gen2B antibodies (2.5 μg i.c.v.) maintained normal hippocampal LTP. (**C**) Similarly, robust LTP was induced when Gen2B mAb (2.5 μg i.c.v.) was co-administered with the extract AD6. Left-hand panels in **B, C** show the time course of LTP. Summary bar charts of LTP magnitude during the last 10 min are displayed in the right-hand panels. Values are mean ± SEM. ^##^p < 0.01, ^###^p < 0.001 compared with pre-HFS, paired *t*-test; ***p < 0.001, ****p < 0.0001, one-way ANOVA followed by Bonferroni’s multiple-comparison tests in **B** and unpaired *t*-test in **C**.

Among the fractions tested in electrophysiological experiments, fractions 15 and 16 significantly inhibited LTP when compared with vehicle-injected control animals (Fig. 2B). Similar to unfractionated Ts21 secretomes (Fig. 1S), ID of one of the active fractions (Fr 15) with the MR-directed mAb Tau5, but not one directed to the extreme CT of tau (Tau46), prevented the disruption of synaptic plasticity (Fig. 3S B). Significantly, co-injection of Fr 15 with the Gen2A mAb, to neutralise tau fragments containing the CT sequence proximal to R’, was also effective in preventing the inhibition of LTP (Fig. 2C).

Taken together these findings implicate MTBR/R’-containing and related fragments in mediating the synaptic plasticity disrupting actions of Ts21 iN secretomes. To test the hypothesis that such tau fragments are sufficient to mediate the synaptotoxicity of tau we investigated the ability of a recombinant human tau preparation to mimic the inhibition of LTP by the secretomes. We chose a tau fragment peptide, tau297-391, which is the dominant species present in AD brain paired helical filaments [16], and which extends beyond the classical MTBR to include the R’ Gen2B epitope sequence. Intriguingly, this peptide robustly inhibited LTP (Fig. 2D).

### Anti-MTBR/R’ tau antibodies abrogate synaptotoxicity of AD brain soluble tau

Having established the ability of antibodies targeting R’ and sequences adjacent to the N- and C-termini of MTBR/R’ to prevent inhibition of LTP by Ts21 extracellular tau fragments, we evaluated the involvement of MTBR-related tau in synaptic plasticity disruption by AD brain tau. To this end we used synaptotoxic tau-containing aqueous brain extracts of two patients diagnosed with sporadic AD (referred to as AD1 and AD6) [17-19]. Western blotting (Fig. 3A) revealed that, apart from full-length tau, Tau5 and the R’-directed Gen2B mAb detected mainly truncated tau fragments, some of which appeared to be recognised by both mAbs.

Consistent with a role for MTBR/R’-containing tau species, co-injection of AD1 extract with either Gen2A or Gen2B mAbs completely restored LTP (Fig. 3B). Furthermore, we discovered the same beneficial effect on LTP of Gen2B mAb when co-injected with AD6 extract (Fig. 3C).

Finally, we wondered if MTBR/R’-containing tau also mediated a very persistent synaptotoxic effect of AD brain tau[19] (Fig. 4A). Remarkably, human tau was detectable in rat hippocampus 3 weeks after a single i.c.v. injection of AD1 extract (Fig. 4B and Fig. 4S). Furthermore, i.c.v. injection of Tau5 mAb just 15 min prior to HFS significantly reversed the LTP deficit in animals injected with AD1 extract 3 weeks earlier (Fig. 4C), consistent with an MR tau-dependence of the persistent LTP inhibition and our previous report using AD6 extract [19]. Importantly, administration of either Gen2A or Gen2B mAbs also rapidly reversed the LTP deficit in this model (Fig. 4D).

**Fig. 4.**
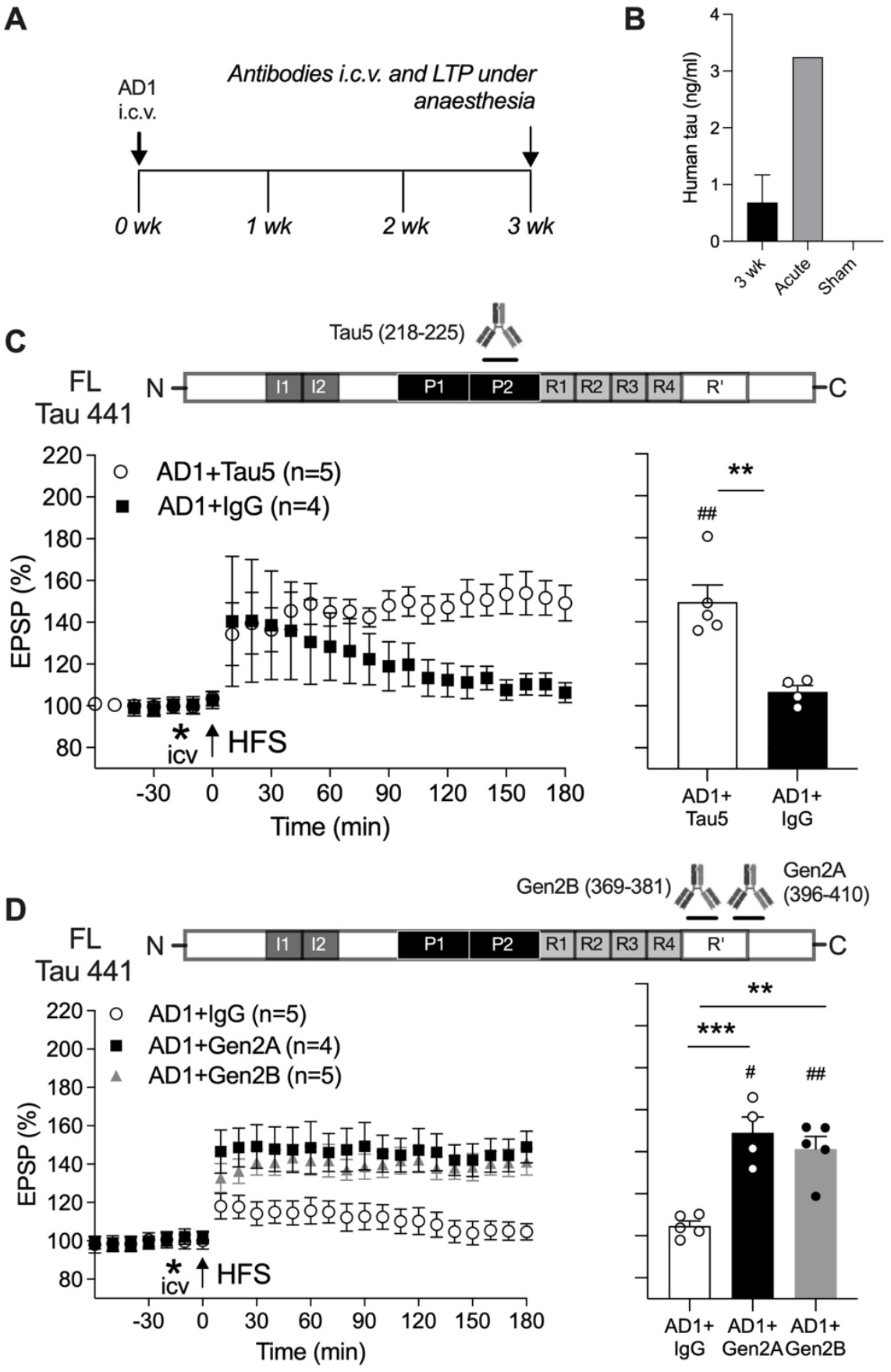
Rapid reversal of the persistent inhibition of LTP by synaptotoxic tau in AD aqueous brain extract by acute injection of anti-tau antibodies. (**A**) Study design. Animals received an i.c.v. injection of AD1 extract under recovery anaesthesia (ketamine and medetomidine, 60 and 0.4 mg/kg, respectively, i.p.), followed 3 weeks later by a single injection with anti-tau or control antibodies under non-recovery urethane anaesthesia. (**B**) Human tau was detected in rat hippocampus 3 weeks (3 wk, n=16 after AD1 i.c.v. injection (LLoQ: 0.03125 ng/ml). For comparison, the levels of human tau detected after acute AD1 (30 min, n=3) with the same volume or sham (n=1) treatment are also shown. (**C**) Whereas LTP was strongly inhibited in animals receiving control mAb IgG (AD1+IgG), robust LTP was induced in rats injected with the mid-region anti-tau mAb Tau5 (AD1+Tau5) (2.5 μg i.c.v., 15 min prior to HFS). (**D**) The same protocol was followed using anti-tau mAbs targeting the MTBR/R’ or MTBR/R’ and the adjacent CT region of tau with Gen2B (AD1+Gen2B) or Gen2A (AD1+Gen2A), respectively. HFS triggered stable LTP in these animals. Left-hand panels in **C, D** show the time course of LTP. Summary bar charts of LTP magnitude during the last 10 min are displayed in the right-hand panels. Values are mean ± SEM. ^#^*p* < 0.05, ^##^*p* < 0.01 compared with pre-HFS, paired *t*-test; \*\**p* < 0.01, \*\*\**p* < 0.001 unpaired *t*-test in (**C**) and one-way ANOVA followed by Bonferroni’s multiple-comparison tests in (**D**).

## Discussion

There is a rapidly growing clinical interest in exploring ways of both treating AD and other tauopathies with MTBR tau-directed therapies and also monitoring disease status with MTBR tau-based fluid biomarkers. Apart from being a major component of tau tangles, MTBR/R’ is also present in a soluble form in AD brain [10, 25] and human CSF [2, 9, 10, 21]. Understanding the involvement of MTBR tau in disease mechanisms should greatly inform these clinical developments. Here, we provide convincing evidence that soluble MTBR/R’-containing tau mediates the synaptotoxicity of patient-derived tau. Either co-injection or ID with two novel MTBR/R’-directed antibodies prevented the acute inhibition of hippocampal LTP by two different AD-relevant preparations: Down syndrome Ts21 iN secretomes and aqueous extracts of AD brain. Furthermore, *in vivo* administration of these agents rapidly reversed the persistent disruption of synaptic plasticity by the brain extracts. Taken together with their potential involvement in the spread of pathology [22], our findings lend new mechanistic support for a focus on soluble MTBR/R’-containing tau species for future therapy and diagnostics.

Our present discovery that two anti-tau antibodies targeting MTBR/R’ and the adjacent CT region, taken together with our previous reports with the MR-directed antibody Tau5 [12, 17-19], abrogated the ability of both of the patient-derived tau preparations to inhibit LTP underlines the importance of tau species containing these regions in mediating synaptotoxicity. In the case of Ts21 iNs, a wide range of putatively extracellular truncated tau species are present in the secretome (Fig. 1S and Hu et al. [12]). In the latter study [12] secretomes from familial AD PS1 L113_I114insT iNs had a similar tau fragment profile but inhibited LTP in an Aß-dependent but tau-independent manner. In the present study, the Ts21 secretomes were found to contain significant amounts of fragments that included both the MR and MTBR/R’ sequences recognised by Tau5 and Gen2B, respectively. Suggestively, these fragments, detected using a novel immunoassay, were markedly reduced by ID with these mAbs (Fig. 1), strongly implicating fragments that encompass both MR and MTBR/R’ as mediators of the inhibition of LTP by Ts21 iN secretomes. The ability of K9JA, targeting both the MTBR and CT, to prevent the inhibition of LTP by Ts21 iN secretomes, unlike Tau46, which targets the extreme CT, is consistent with this conclusion. However, the efficiency of ID with Tau46 was relatively weak. In the case of the AD aqueous extracts it is not possible differentiate if synaptotoxicity is mediated by full-length tau or one or more MTBR/R’-containing fragments, or whether the active species are derived from within neurons or extracellularly. Nonetheless, data from AD brain extracts confirm the requirement of the MR and MTBR/R’ domains for synaptotoxic activity. In view of the protective activity of Gen2A (Fig. 1,3; Fig. 2S), the rogue tau species also likely include peptides incorporating the CT sequence flanking MTBR/R’. The ability of recombinant tau297-391 to replicate the synaptic plasticity disrupting action of patient-derived tau supports the hypothesis that peptides containing a core MTBR/R’ tau sequence are necessary and sufficient to mediate the synaptotoxicity of tau. We and others previously reported that recombinant preparations of full-length tau needed to be pre-aggregated to inhibit LTP [6, 17-19] whereas tau297-391 appears to be a very potent synaptotoxin without deliberate aggregation. Although highly unlikely in the brief time course of the present experiments, we cannot rule out the possibility that this aggregation-prone tau species may either contain low levels of oligomers or oligomerise quickly after injection of monomers into live rat brain. Similarly, although it is feasible that fragments containing tau peptides spanning from the MR to just CT of MTBR/R’, it is also possible that at least two small fragments of tau, one MR and the other MTBR/R’-containing, are acting in concert to cause synaptic plasticity impairment. Interestingly, the most potent SEC fractions of Ts21 iN secretomes contain both MR and MTBR-containing fragments. Because AD aqueous brain extracts contain a very wide range of intracellular and extracellular post-translationally-modified tau species it is likely that the nature of the synaptotoxic culprits in these preparations is even more complex than those in the Ts21 iN secretomes.

We confirmed and extended our previous reports that patient-derived synaptotoxic tau-containing brain extracts potently inhibit LTP for several weeks *in vivo* [18, 19]. The rapid reversal of the deficit by the two anti-MTBR/R’ antibodies using 15-min i.c.v. treatment, 3 weeks after AD1 extract exposure, indicates the prolonged presence of readily accessible synaptotoxic AD tau. Consistent with this hypothesis, human tau was detectible in the hippocampus of these animals at this time (Fig. 4). Given that only a brief exposure was needed for the antibodies to exert their protective action, it is likely that they directly bound and thereby neutralised the synaptotoxic MTBR/R’-containing tau species.

In conclusion, our discovery that MTBR/R’-containing and related fragments are potent mediators of the synaptic plasticity-disrupting actions of patient-derived soluble synaptotoxic tau provides a biological basis for interpreting ongoing biomarker-based clinical trials in AD immunotherapy.

## Supplementary information

Below is the link to supplementary material. Supplementary material

## Author Contributions

Conceptualization: I.K., M.J.R., D.M.W. and F.J.L. Data collection: I.K., T.O., N.W.H., E.C., T.C., G.T.C. and I.D. Supervision: M.J.R, F.J.L., K.M. and D.M.W. Writing original draft: I.K. and M.J.R. All authors reviewed the manuscript.

## Data availability

The original data are available upon reasonable request.

## Acknowledgements

We thank Dr. Matthew Frosch of MGH for providing AD brain tissue.

## Declarations Ethics approval

Human brain tissue was used in accordance with Trinity College Dublin Faculty of Health Science Ethics Committee (approval 16014) and Partners Institutional Review Board (protocol, Walsh BWH 2011) guidelines. This research was carried out in accordance with the UK Code of Practice for the Use of Human Stem Cell Lines. All animal experiments were conducted in accordance with ARRIVE (Animal Research: Reporting of *In Vivo* Experiments) guidelines under the approval of Trinity College Dublin local animal research ethics committee and the Health Products Regulatory Authority in Ireland.

