## Supplementary material for "Synaptotoxic effects of extracellular tau are mediated by its microtubule-binding region"

(1) Figures 1-5S

(2) Tables 1-2S

(3) Supplementary Materials and Methods

(4) Supplementary References

### Supplementary Figures

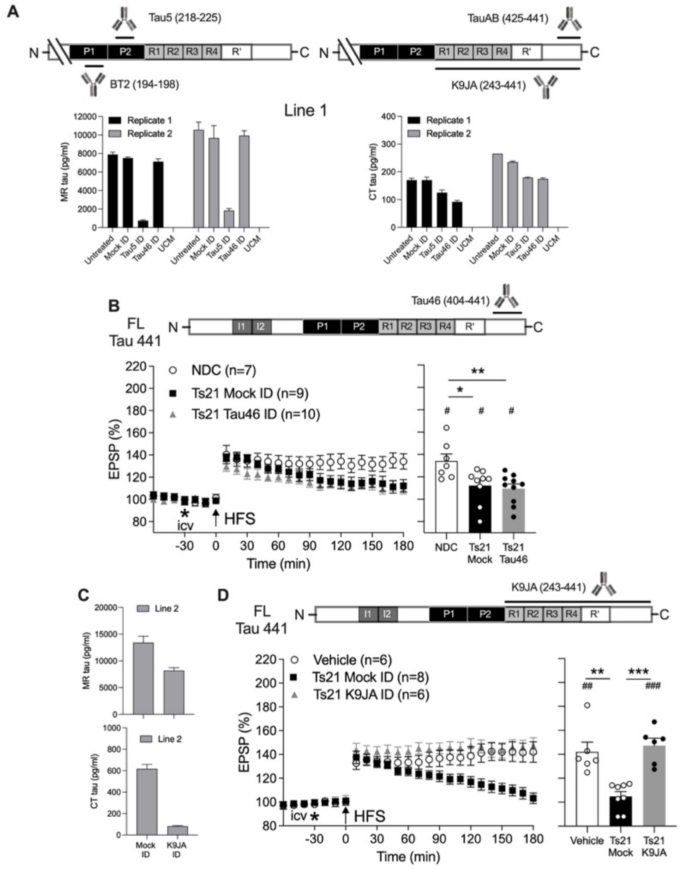


Figure 1S. **Effect of anti-tau antibodies directed to the MTBR/CT and extreme CT on the synaptotoxicity of patient-derived extracellular tau** (**A**) Insets show antibodies used to capture/detect mid-region (MR) and C-terminal (CT)-containing tau fragments in sandwich ELISAs. Effect of ID with MR-directed Tau5 and extreme CT-directed Tau46 anti-tau mAbs on levels of MR and CT-containing tau fragments. Secretomes from line 1 of two lines of Ts21 iNs or unconditioned media (UCM) were assayed twice using Tau5-BT2 and K9JA-TauAB sandwich ELISAs. Tau5 ID reduced MR-containing tau fragments by ~95% and CT pool by ~30%. Tau46 ID reduced the CT pool by ~30-40%, with little detectible effect on MR tau. In contrast to Tau46 (**B**), K9JA ID lowered both CT and, to a lesser extent, MR tau levels (**C**) and prevented inhibition of hippocampal LTP under urethane anaesthesia (**D**). Left-hand panel shows the time course of LTP. Summary bar chart of LTP magnitude during the last 10 min is in the right-hand panel. Values are mean ± SEM. $^{\#}p < 0.05$, $^{\#\#\#\#}p < 0.0001$ compared with pre-HFS, paired *t*-test; *p < 0.05, **p < 0.01, ***p < 0.001, one-way ANOVA followed by Bonferroni's multiple-comparison tests.

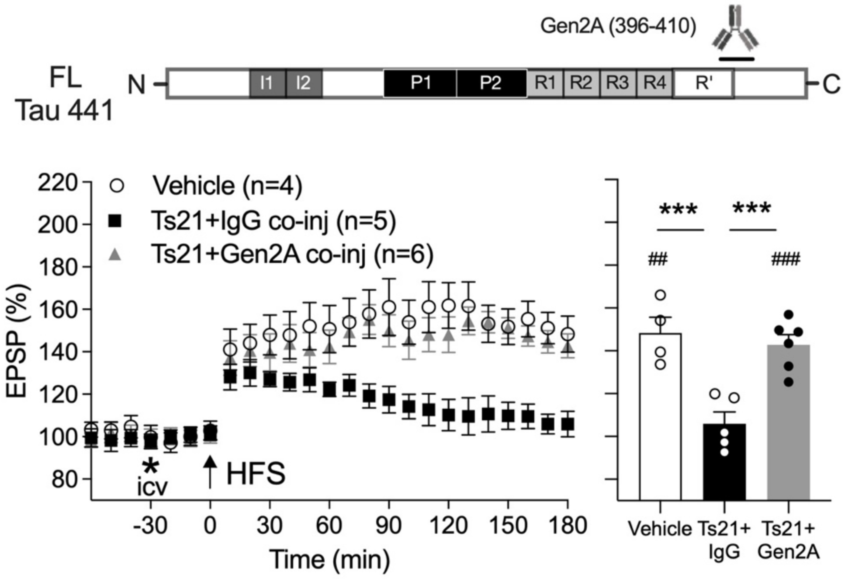


Figure 2S. **Co-injection of an antibody directed C-terminal of the R' domain (Gen2A) abrogates Ts21 iN secretome-induced LTP deficit.** I.c.v. injection of Gen2A (2.5 µg)-containing Ts21 iN secretome failed to inhibit LTP, unlike co-injection of an isotype control IgG (2.5 µg). Left-hand panel shows the time course of LTP. Summary bar chart of LTP magnitude during the last 10 min is in the right-hand panel. Values are mean ± SEM. $^{\#\#}p < 0.01$, $^{\#\#\#}p < 0.001$ compared with pre-HFS, paired *t*-test; ***p < 0.001, one-way ANOVA followed by Bonferroni's multiple-comparison tests.

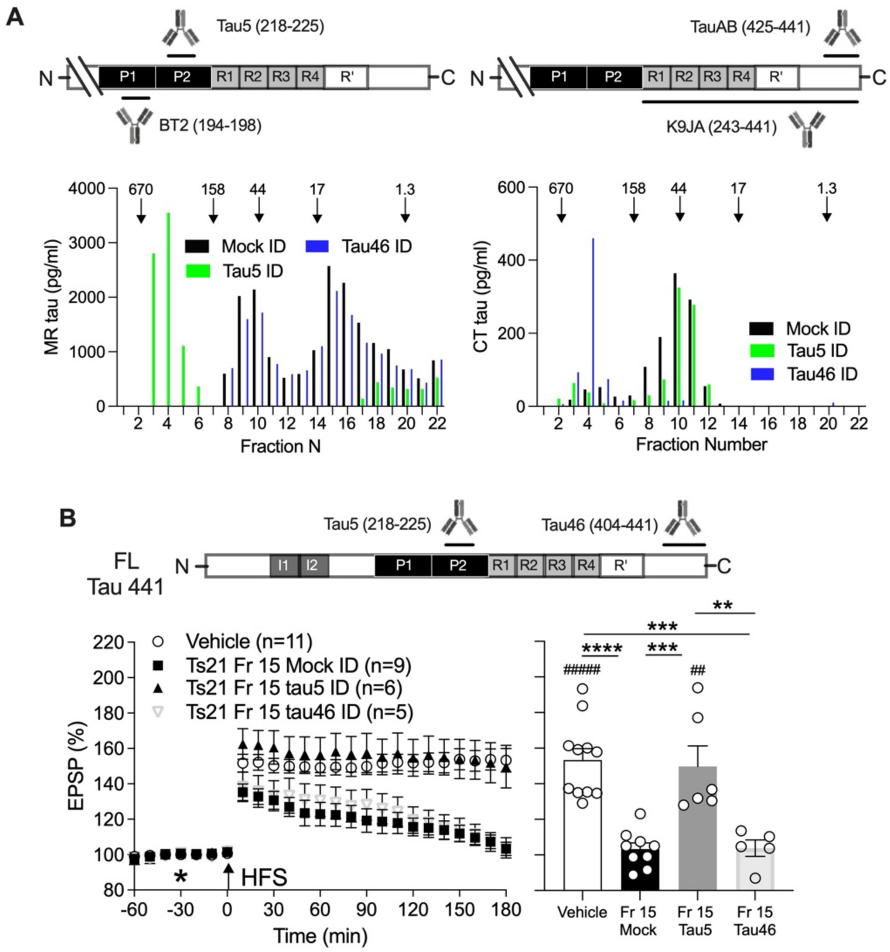


Figure 3S. **Abrogation of the inhibition of LTP by active SEC fraction of Ts21 iN secretome by immunodepletion with mid-region-directed antibody** (**A**) Effect of ID with mid-region (MR) directed Tau5 and C-terminally (CT) directed Tau46 anti-tau mAbs on MR and CT tau fragments. The concentration of MR and CT-containing tau in the different SEC fractions were assayed using Tau5-BT2 and K9JA-TauAB sandwich ELISAs, respectively. The elution of globular protein standards (the molecular weight of which are given in kDa) is indicated by downward pointing arrows on the top of the chromatogram. Whereas active Ts21 Fr 15 Tau5 ID prevented inhibition of LTP (**B**), Tau46 ID did not (**C**). Left-hand panel shows the time course of LTP. Summary bar chart of LTP magnitude during the last 10 min is in the right-hand panel. Values are mean ± SEM. $^{\#\#}p < 0.01$, $^{\#\#\#\#}p < 0.0001$ compared with pre-HFS, paired *t*-test; $^{**}p < 0.01$, $^{***}p < 0.001$, $^{****}p < 0.0001$, one-way ANOVA followed by Bonferroni's multiple-comparison tests.

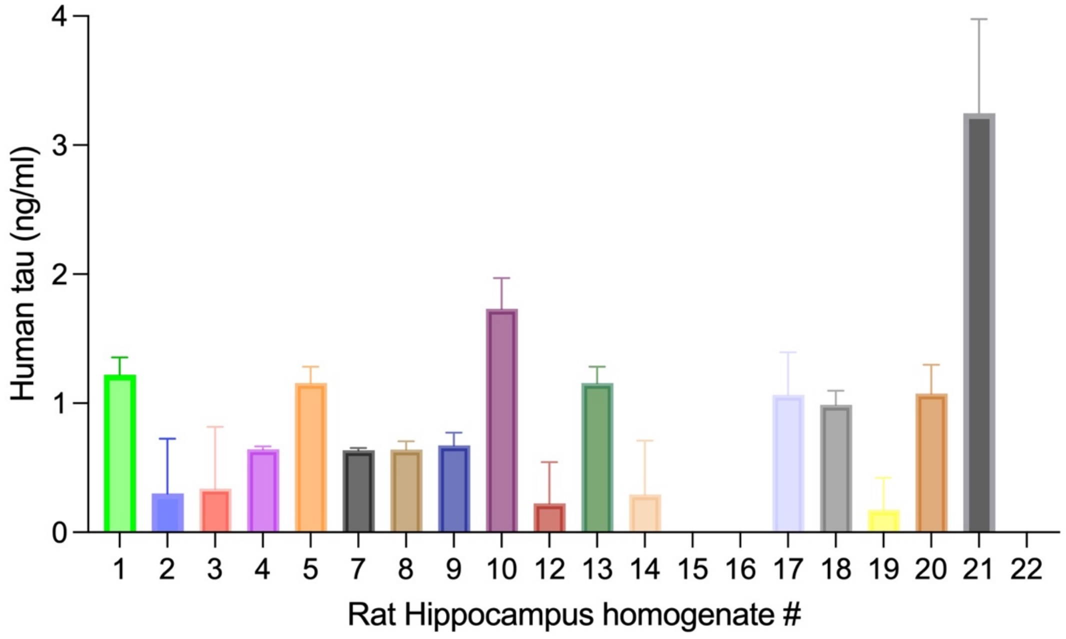


Figure 4S. **Concentration of human tau in individual rat hippocampal homogenates after intracerebroventricular injection of Alzheimer's disease soluble brain extract** Three weeks after animals # 1-20 received a single i.c.v. injection of AD1 aqueous extract (either Mock ID, # 1-10, or AW7 ID, # 12-20) under recovery anaesthesia their brains were removed under non-recovery anaesthesia and the concentration of human tau assayed in hippocampal homogenates. Homogenate # 21 is a pooled sample from three rats acutely injected i.c.v. with the same volume of AD1 Mock ID extract 30 min previously. Animal # 22 received a single intrahippocampal acute sham injection. LLoQ was 0.03125 ng/ml.

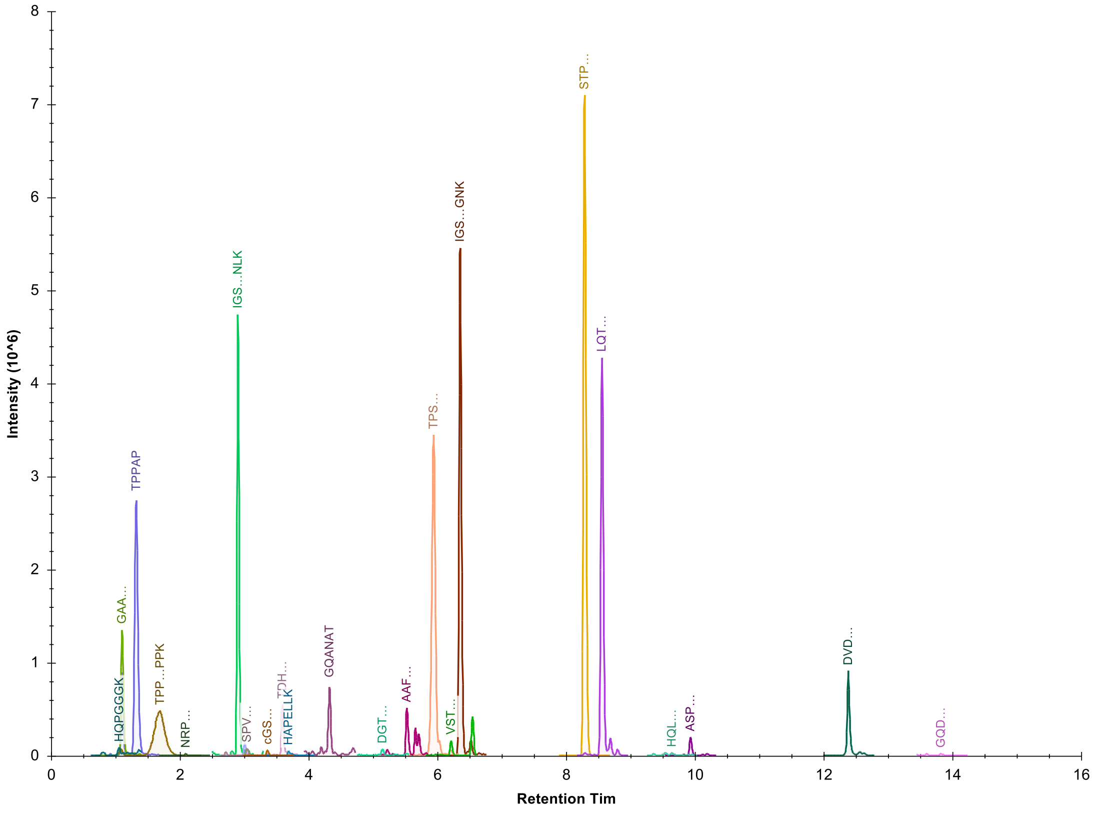


Figure 5S **Example of peptide chromatography- Pooled quality control (QC) sample**

### Supplementary Tables

Table 1S **Primary antibodies and their antigens, dilutions and sources.**

| Antibody | Clonality | Antigen/Epitope | Dilution | Source | Reference |
|---|---|---|---|---|---|
| 46-4 | Monoclonal | Anti-HIV | 1:100 (IP) | ATCC | Wang et al. [13] |
| BT2 | Monoclonal | Tau 194-198 | 2.5 μg/ml (ELISA) | ThermoFisher | Mercken et al. [5] |
| Gen2A | Monoclonal | Tau 396-410 | 1:100 (IP);<br>2.5 μg (i.c.v.) | Gen2 Neuroscience | In-house antibody |
| Gen2B | Monoclonal | Tau 369-381 | 1:100 (IP);<br>4 μg/ml (MSD);<br>1:1000 (WB);<br>2.5 μg (i.c.v.) | Gen2 Neuroscience | In-house antibody |
| IgG1κ | Monoclonal | N/A | 2.5 μg (i.c.v.) | BioLegend | Ondrejcak et al. [8] |
| IgG rabbit | Monoclonal | N/A | 1:100 (IP);<br>2.5 μg (i.c.v.) | eBioscience | Bode et al. [1] |
| K9JA | Polyclonal | Tau 243-441 | 1:100 (IP);<br>2.5 μg/ml (ELISA) | Dako | Wang et al. [13] |
| Tau46 | Monoclonal | Tau 404-441 | 1:100 (IP) | BioLegend | Meredith et al. [6] |
| Tau5 | Monoclonal | Tau 218-225 | 1:100 (IP);<br>2.5 μg/ml (ELISA);<br>2.5 μg (i.c.v.) | BioLegend | Porzig et al. [9] |
| | | | 2 μg/ml (MSD); 1:500 (WB) | ThermoFisher | |
| TauAB | Monoclonal | Tau 425-441 | 2.5 μg/ml (ELISA) | MedImmune | Hu et al. [2] |

Table 2S **Table of transitions used in the targeted assay**

| Peptide Sequence | Precursor Mz | Precursor Charge | Collision Energy | Product Mz | Product Charge | Fragment Ion |
|---|---|---|---|---|---|---|
| STPTAEDVTAPLVDEGAPGK | 652.325 | 3 | 22.2 | 982.5204 | 1 | y10 |
| STPTAEDVTAPLVDEGAPGK | 652.325 | 3 | 22.2 | 673.3151 | 1 | y7 |
| STPTAEDVTAPLVDEGAPGK | 652.325 | 3 | 22.2 | 491.7638 | 2 | y10 |
| HAPELLK | 404.2398 | 2 | 13.9 | 670.4134 | 1 | y6 |
| HAPELLK | 404.2398 | 2 | 13.9 | 599.3763 | 1 | y5 |
| HAPELLK | 404.2398 | 2 | 13.9 | 435.1987 | 1 | b4 |
| HQLLGDLHQEGPPLK | 561.3055 | 3 | 18.9 | 768.425 | 1 | y7 |
| HQLLGDLHQEGPPLK | 561.3055 | 3 | 18.9 | 453.2456 | 2 | y8 |
| HQLLGDLHQEGPPLK | 561.3055 | 3 | 18.9 | 664.3413 | 1 | b6 |
| DVDESSPQDSPPSK | 496.5547 | 3 | 16.5 | 515.2824 | 1 | y5 |
| DVDESSPQDSPPSK | 496.5547 | 3 | 16.5 | 428.2504 | 1 | y4 |
| DVDESSPQDSPPSK | 496.5547 | 3 | 16.5 | 428.214 | 2 | y8 |
| ASPAQDGRPPQTAAR | 761.8897 | 2 | 27.1 | 740.405 | 1 | y7 |
| ASPAQDGRPPQTAAR | 761.8897 | 2 | 27.1 | 682.8551 | 2 | y13 |
| ASPAQDGRPPQTAAR | 761.8897 | 2 | 27.1 | 477.2674 | 2 | y9 |
| VSTEIPASEPDGPSVGR | 566.6162 | 3 | 19.1 | 784.3948 | 1 | y8 |
| VSTEIPASEPDGPSVGR | 566.6162 | 3 | 19.1 | 572.3151 | 1 | y6 |
| VSTEIPASEPDGPSVGR | 566.6162 | 3 | 19.1 | 392.701 | 2 | y8 |
| GQDAPLEFTFHVEITPNVQK | 757.3866 | 3 | 25.9 | 686.3832 | 1 | y6 |
| GQDAPLEFTFHVEITPNVQK | 757.3866 | 3 | 25.9 | 585.3355 | 1 | y5 |
| GQDAPLEFTFHVEITPNVQK | 757.3866 | 3 | 25.9 | 950.0042 | 2 | y16 |
| AAFPGAPGEGPEAR | 442.8861 | 3 | 14.6 | 715.3369 | 1 | y7 |
| AAFPGAPGEGPEAR | 442.8861 | 3 | 14.6 | 529.2729 | 1 | y5 |
| AAFPGAPGEGPEAR | 442.8861 | 3 | 14.6 | 472.2514 | 1 | y4 |
| DGTGSDDK | 397.6618 | 2 | 13.6 | 679.2893 | 1 | y7 |
| DGTGSDDK | 397.6618 | 2 | 13.6 | 521.2202 | 1 | y5 |
| DGTGSDDK | 397.6618 | 2 | 13.6 | 464.1987 | 1 | y4 |
| NRPCLSPK | 486.2582 | 2 | 16.9 | 528.2347 | 1 | b4 |
| NRPCLSPK | 486.2582 | 2 | 16.9 | 641.3188 | 1 | b5 |
| NRPCLSPK | 486.2582 | 2 | 16.9 | 728.3508 | 1 | b6 |
| GAAPPGQK | 363.2007 | 2 | 12.4 | 597.3355 | 1 | y6 |
| GAAPPGQK | 363.2007 | 2 | 12.4 | 526.2984 | 1 | y5 |
| GAAPPGQK | 363.2007 | 2 | 12.4 | 263.6528 | 2 | y5 |
| GQANATR | 359.1856 | 2 | 12.2 | 660.3424 | 1 | y6 |
| GQANATR | 359.1856 | 2 | 12.2 | 532.2838 | 1 | y5 |
| GQANATR | 359.1856 | 2 | 12.2 | 461.2467 | 1 | y4 |
| TPPAPK | 305.6816 | 2 | 10.2 | 412.2554 | 1 | y4 |
| TPPAPK | 305.6816 | 2 | 10.2 | 255.1577 | 2 | y5 |
| TPPAPK | 305.6816 | 2 | 10.2 | 206.6314 | 2 | y4 |
| TPPSSGEPPK | 498.7535 | 2 | 17.4 | 798.3992 | 1 | y8 |

| | | | | | | |
|---|---|---|---|---|---|---|
| TPPSSGEPPK | 498.7535 | 2 | 17.4 | 448.2296 | 2 | y9 |
| TPPSSGEPPK | 498.7535 | 2 | 17.4 | 399.7032 | 2 | y8 |
| SGYSSPGSPGTPGSR | 697.3208 | 2 | 24.7 | 912.4534 | 1 | y10 |
| SGYSSPGSPGTPGSR | 697.3208 | 2 | 24.7 | 671.3471 | 1 | y7 |
| SGYSSPGSPGTPGSR | 697.3208 | 2 | 24.7 | 456.7303 | 2 | y10 |
| TPSLPTPPTR | 533.7982 | 2 | 18.7 | 868.4887 | 1 | y8 |
| TPSLPTPPTR | 533.7982 | 2 | 18.7 | 668.3726 | 1 | y6 |
| TPSLPTPPTR | 533.7982 | 2 | 18.7 | 286.1636 | 2 | y5 |
| LQTAPVPMPDLK | 655.3629 | 2 | 23.2 | 1068.5758 | 1 | y10 |
| LQTAPVPMPDLK | 655.3629 | 2 | 23.2 | 896.491 | 1 | y8 |
| LQTAPVPMPDLK | 655.3629 | 2 | 23.2 | 700.3698 | 1 | y6 |
| IGSTENLK | 431.2375 | 2 | 14.9 | 748.3836 | 1 | y7 |
| IGSTENLK | 431.2375 | 2 | 14.9 | 691.3621 | 1 | y6 |
| IGSTENLK | 431.2375 | 2 | 14.9 | 604.3301 | 1 | y5 |
| HQPGGGK | 340.6774 | 2 | 11.5 | 543.2885 | 1 | y6 |
| HQPGGGK | 340.6774 | 2 | 11.5 | 415.23 | 1 | y5 |
| HQPGGGK | 340.6774 | 2 | 11.5 | 318.1772 | 1 | y4 |
| CGSLGNIHHKPGGGQVEVK | 658.6707 | 3 | 22.4 | 870.468 | 1 | y9 |
| CGSLGNIHHKPGGGQVEVK | 658.6707 | 3 | 22.4 | 878.9763 | 2 | y17 |
| CGSLGNIHHKPGGGQVEVK | 658.6707 | 3 | 22.4 | 778.9182 | 2 | y15 |
| IGSLDNITHVPGGGNK | 526.946 | 3 | 17.6 | 529.2729 | 1 | y6 |
| IGSLDNITHVPGGGNK | 526.946 | 3 | 17.6 | 733.3733 | 2 | y15 |
| IGSLDNITHVPGGGNK | 526.946 | 3 | 17.6 | 704.8626 | 2 | y14 |
| TDHGAEIVYK | 378.1926 | 3 | 12.3 | 522.3286 | 1 | y4 |
| TDHGAEIVYK | 378.1926 | 3 | 12.3 | 482.1994 | 1 | b5 |
| TDHGAEIVYK | 378.1926 | 3 | 12.3 | 611.242 | 1 | b6 |
| SPVVSGDTSPR | 551.2804 | 2 | 19.3 | 917.4687 | 1 | y9 |
| SPVVSGDTSPR | 551.2804 | 2 | 19.3 | 818.4003 | 1 | y8 |
| SPVVSGDTSPR | 551.2804 | 2 | 19.3 | 719.3319 | 1 | y7 |

### Supplementary Materials and Methods

#### Secretome collection, clarification and dialysis

Human pluripotent stem cells were maintained on Geltrex in Essential 8 media (both ThermoFisher). Directed differentiation of iPSCs to cerebral cortex neural progenitor cells and neurons was performed as previously described [10-12]. Secretomes of differentiated neurons was collected between day 70 and day 80 post-neural induction at 48 h intervals, aliquoted and stored in protein lo-bind tubes at -80°C.

Majority of secretome preparations (Fig 1B,D; Fig 1S, Fig 2S A-C; Fig 3S) were cleared of cell debris and extracellular vesicles by centrifugation. First, secretome was centrifuged at 200 × g and 4°C for 10 min. Then, the upper 97% was recovered (supernatant 1; S1) and centrifuged at 2,000 × g and 4°C for 10 min. Next, the upper 97% of this was recovered (S2) and centrifuged at 10,000 × g and 4°C for 30 min. Finally, the upper 97% was recovered (S3) and centrifuged at 100,000 × g and 4°C for 70 min.

All secretomes were dialyzed (using Slide-A-Lyzer™ G2 Dialysis Cassettes, 2K MWCO, ThermoFisher) against artificial cerebrospinal fluid (aCSF; 124 mM NaCl, 2.8 mM KCl, 1.25 mM NaH2PO4, 26 mM NaHCO3) to remove bioactive small molecules. Dialysis was performed at 4°C against a 100-fold excess of aCSF with buffer changed three times over a 48-h period. Dialyzed secretome was either frozen at -80°C or used immediately for further biochemical manipulations.

#### Preparation of soluble human brain extracts

Both Alzheimer's disease cases met current post-mortem diagnostic criteria (see Table 1 in Ondrejcak *et al*, [8]). Brain sample from AD1 was obtained from the Massachusetts Alzheimer's Disease Research Center Neuropathology Core at Massachusetts General Hospital. AD6 brain was from the Banner Health brain bank. Aqueous brain extracts were prepared from cortical grey matter by clarification and dialysis (2K molecular weight cutoff) of 20% (w/v) homogenates, as described in our previous publications [7, 8].

#### Size exclusion chromatography (SEC)

Twelve milliliter aliquots of ultracentrifuged and dialyzed secretomes were concentrated 10-fold (to 1.2 mL) using Amicon Ultra-15 3 kDa centrifugal filters (Millipore, Billerica, MA) at 4ºC. Immediately thereafter, 1 mL of concentrate was chromatographed on tandem Superdex 200 Increase - Superdex 75 10/300 GL (GE, Marlborough, MA) columns eluted in 50 mM ammonium bicarbonate (pH 8.5) at a flow rate of 0.5 mL/min using Pharmacia FPLC system (Amersham Life Sciences, Uppsala, Sweden). One milliliter fractions were collected. SEC was repeated twice using both Ts21 line 1 and line 2. A 900 µL of each fraction was lyophilized and the remaining volume aliquoted and stored at -80ºC pending analyses using tau immunoassays and electrophysiology. Since it was necessary to concentrate conditioned media prior to SEC, we were careful to confirm that the concentration step did not remove synaptotoxic activity. Similar to unconcentrated media, concentrated Ts21 secretomes inhibited LTP *in vivo*, and this inhibition was prevented by Tau5 immunodepletion (Ts21 Mock ID, n=4, vs Ts21 Tau5 ID, n=4, p=0.012, data not shown).

#### Antibodies

The antibodies used and their sources are described in Table 1S.

#### Compounds

$Tau_{297-391}$, also known as dGAE, the truncated recombinant human tau fragment encompassing the core MTBR/R'- region present in tau fibrils, was kindly provided by StressMarq Biosciences Inc. (Victoria, Canada) in its monomer form.

#### Tau enzyme-linked immunosorbent assays (ELISAs)

Assays for mid-region (MR) and C-terminal (CT) tau in Ts21 secretomes were performed using the same procedure but with specific combinations of capture and detection antibodies [3]. For all assays, the capture antibody was coated at 2.5 µg/mL in Tris-buffered saline (TBS) for 1 h at 37°C and 300 rpm agitation. Plates were then washed three times with 100 µL TBS with Tween 20 (TBST) prior to blocking in 100 µL TBS containing 3% BSA for 2 h at RT and 300 rpm. Plates were washed three times with 100 µL TBST before 25 µL of samples (diluted 1:1 in TBS containing 1% BSA) and standards were applied in triplicate and agitated for 16 h at 4°C. Importantly, the same calibration standard (recombinant human tau441) was used for all assays, thus enabling comparison of concentrations detected by different assays. The following day, 25 µL alkaline phosphatase conjugated detection antibodies diluted 1:250 in TBST containing 1% BSA were added directly to the plates without washing and incubated for 1 h at RT and 300 rpm. Finally, plates were washed three times with 100 µL TBST before 50 µL Tropix Sapphire II (Applied Biosystems) detection reagent was added and incubated for 30 min at RT and 300 rpm. Standard curves were fitted to a five-parameter logistic function with 1/Y2 weighting using MasterPlex ReaderFit (MiraiBio).

#### Custom MSD (Meso Scale Diagnostics) assay for tau containing the MTBR/R' domain

All solutions were prepared in PBS, supplemented with 5% BSA for blocking solution, 1% BSA for detection antibody solution, or 0.05% Tween-20 for wash solution. Plates were coated overnight with 4 µg/mL monoclonal rabbit anti-tau (aa369-375) capture antibody (Gen2B), followed by blocking for 1 h. Conditioned media samples or recombinant full-length monomeric tau (rPeptide, T-1001) were incubated for 3 h. For detection, plates were incubated overnight with 2 µg/mL monoclonal Tau5 mouse anti-tau (aa218-225) detection antibody (ThermoFisher Scientific, AHB0042) and detected with 1 µg/mL goat anti-mouse SULFO-TAG secondary antibody (MSD, R32AC) for 1 h. Signal was generated using MSD Gold Read Buffer B and detected by MESO QuickPlex SQ 120MM Imager (MSD). Concentrations of MTBR/R'-containing tau in conditioned media samples were calculated with reference to the standard curve using MSD Discovery Workbench software.

#### Rat hippocampal tissue lysate preparation

After euthanasia, hippocampi were dissected and stored frozen. Immediately prior to analysis, tissue was thawed and homogenized in Pierce™ RIPA buffer (ThermoFisher Scientific, #89900, one part of tissue to 7 parts of lysis buffer) containing protease and phosphatase inhibitor cocktail (Sigma-Aldrich, #P8849, 1mM final concentration) to ensure protein integrity. Tissue and cell debris was removed by centrifugation (10000 x g for 5 min at 4°C). To measure human tau in rat hippocampal brain homogenates (Fig 4B), total human tau ELISA kit (Invitrogen, #KHB0041) was used in accordance with manufacturer's instructions.

#### Immunodepletion (ID)

Ts21 secretomes underwent 1-3 rounds of 12-16 h ID by incubation with Protein A Sepharose beads (PAS; 10 µL, ThermoFisher) at 4°C with either (i) anti-tau antibodies (Tau5, Tau46, K9JA, Gen2A or Gen2B) and or (ii) control antibodies (46-4 or rabbit IgG) or purified pre-immune serum. In each case, samples were cleared of beads and then both the ID and 'mock' ID samples were incubated with PAS alone to remove previously unbound IgG. ID supernatants were stored at -80°C.
To rule out possible involvement of amyloid beta, in some cases, we used Ts21 secretomes (Fig 1S B) and AD1 extract (Fig 3A; Fig 4B, D; Fig 4S) that had been previously subjected to ID treatment with the polyclonal anti-amyloid beta antibody AW7 [2, 8].

#### Western immunoblot analysis

Aqueous extracts from the brains of two people with Alzheimer's disease were analysed for tau protein by western immunoblotting. Samples were run on 4–15% Criterion TGX Stain-Free Protein Gels (Bio-Rad) and proteins transferred to Trans-Blot Turbo Midi 0.2 µm PVDF membranes (Bio-Rad). Membranes were blocked with 5% BSA in TBS/0.1% Tween (TBS-T) for 1 h at RT before an overnight incubation with primary antibodies (monoclonal Gen2B rabbit anti-tau IgG, 1:1000; monoclonal Tau5 mouse anti-tau IgG, 1:500, ThermoFisher Scientific AHB0042), in blocking solution at RT. For detection, membranes were washed in TBS-T following primary antibody incubation and then incubated with fluorescent secondary antibodies (goat anti-rabbit-647 IgG, 1:2000, ThermoFisher Scientific A21244; goat anti-mouse-568 IgG, 1:2000 ThermoFisher Scientific A11031), in blocking solution for 2 h at RT, washed in TBS-T and imaged in the 647 and 568 channels on a ChemiDoc Imaging System (Bio-Rad).

#### Surgery and electrophysiology

Rats were group-housed, unless otherwise stated. Food and water were available *ad libitum* with a 12 h light/dark cycle. For most experiments we used 3-5-month-old male Lister Hooded rats. In the study examining the effect of Gen2A ID (Fig 1G) we employed older (6–7-month-old) Wistar Han rats. Control experiments were interleaved randomly throughout, and no animals were excluded. Totally 274 rats were used in this study.

To inject samples and test synaptic plasticity, the animals were anaesthetized with urethane (1.6 g/kg, i.p.) and core body temperature was maintained at 37.5 ± 0.5 °C. An intracerebroventricular (i.c.v) stainless steel guide cannula (22 gauge, 0.7 mm outer diameter, length 13 mm) was implanted above the right lateral ventricle (coordinates, 0.5 mm posterior to bregma and 1.2 mm right of midline, depth 4 mm) before the electrodes were implanted ipsilaterally. Teflon-coated tungsten wire (external diameter 75 µm bipolar or 112 µm monopolar) electrodes were positioned in the stratum radiatum of area CA1. The electrodes were optimally located using a combination of physiological and stereotactic indicators (3.8 mm posterior to bregma and 2.5 mm lateral to midline, and 4.6 mm posterior to bregma and 3.8 mm lateral to midline for recording and stimulating electrodes, respectively). Screw electrodes located over the contralateral cortex were used as reference and earth. To inject samples acutely, a Hamilton syringe was connected to the internal cannula (28 gauge, 0.36 mm outer diameter). The injector was removed 1 min post-injection and a stainless-steel plug was inserted.

To investigate the persistence of the effects of AD1 brain extract, the i.c.v. injections were administered under recovery anesthesia using a mixture of ketamine and medetomidine (60 and 0.4 mg/kg, respectively, i.p.), and the cannula was then removed. Afterwards, rats were housed individually in their home cages. Subsequently, electrophysiological recordings were conducted under non-recovery urethane anesthesia, as described below, 21 d later. At the end of experiment both hippocampi were dissected, homogenized and frozen at -80°C.

To study synaptic plasticity, field excitatory postsynaptic potentials (EPSPs) were evoked and recorded in the stratum radiatum under urethane anaesthesia. Single square-wave pulse (0.2 ms duration) was applied every 30 s and at intensity that triggered a 50% maximum EPSP response. To induce LTP a 200 Hz high-frequency stimulation (HFS) protocol consisting of one set of 10 trains of 20 pulses (inter-train interval of 2 s) at test intensity was applied. The magnitude of control LTP varied considerably over the period of carrying out these experiments. To minimize the possible confounding effect of such variation in control LTP, the experiments in each study were interleaved.

The timing (15-30 min and 3 weeks prior to HFS, in an acute and delayed models, respectively) and doses/volumes of injection of antibodies (2.5 μg per i.c.v. injection), Ts21 secretomes (15-20 μL per i.c.v. injection), and patient-derived brain aqueous extracts (10 μL per i.c.v. injection) were based on pilot experiments and our previous experience with these materials and models [2, 7, 8].

For electrophysiological experiments, values are presented as the mean ± SEM of pre-HFS baseline EPSP amplitude over a 30 min period. The magnitude of LTP was measured at 3 h post-HFS and expressed as the mean ± SEM % baseline. For graphing purposes, EPSP amplitude measurements were grouped into 10 min (average of 20 sweeps).

#### Mass spectrometry

##### Assay Development

The full sequence of the tau protein was obtained from UNIPROT (August 2023), and unique tryptic peptides were identified. These peptides were subsequently imported into Skyline (MacCoss Lab Software, 23.1.0.268) [4], where, in conjunction with Prosit (www.proteometools.org), an *in silico* spectral library was generated for each peptide across different precursor charge states (+2, +3, and +4). A normalised collision energy (NCE) of 31, which was experimentally established for this instrument, was utilised to create the spectral library. The fragment selection for the specified charge states was refined by filtering to retain the 12 most intense fragments predicted by the *in silico* spectral library. This process facilitated the exportation of multiple acquisition methods to MassLynx (Waters), each configured with a minimum dwell time per transition of 10 μs. Chromatographic conditions and injection volumes are described in subsequent sections. A tryptic peptide digest of human recombinant tau352 with a C-terminal His tag (Abcam, ab316441) was utilised for method development, in addition to the pooled quality control (QC) sample (described in the Pre-analysis Sample Preparation section and Figure 5S). Following data acquisition (Xevo TQ-XS Triple Quadrupole Mass Spectrometer and ACQUITY UPLC I-Class PLUS System, Waters), raw data were imported into Skyline for evaluation, and peptide identity was confirmed through spectral library matching (dot product score cut-off of 0.75). The selection of the predominant precursor charges and fragments was based on the intensity and chromatographic performance of the pooled QC sample. This was followed by an additional round of method export, focusing on collision energy optimisation for selected precursors and fragments (see Table 2S). Utilising the experimental data, a final method with three fragments per peptide was developed for the scheduled acquisition of each peptide within a 0.8-minute window to minimise overlap and enhance sensitivity.

#### Sample Preparation and Digestion Protocol

Lyophilised fractions were re-suspended in 900 µL of $dH_20$ and a 450 µL aliquot from each fraction was transferred into 1.5 mL microtubes and subjected to lyophilisation overnight again. Subsequently, the dried samples were resuspended in 20 µL of digestion buffer (6M urea, 2M thiourea, and 2% ASB-14 in 200 mM Tris-HCl, pH 8.0) and incubated at room temperature for 1 h with agitation to facilitate protein denaturation. Disulfide bond reduction was achieved through the addition of 3 µL of 169 mM dithiothreitol (DTT) and incubation for 1 h at room temperature with shaking. Alkylation was performed by adding 6 µL 169 mM iodoacetamide (IAA) and incubating for 45 min in the dark at room temperature. To facilitate efficient trypsin digestion, the mixture was diluted with 165 µL of Milli-Q water to reduce the urea concentration below 1M. Trypsin Gold (Promega, V5280) was prepared at a concentration of 0.1 µg/µL in 200 mM Tris-HCl (pH 8.0). Digestion was initiated by adding 10 µL trypsin solution (1 µg) to each sample, followed by brief vortexing. The samples were then subjected to digestion for 16 h at 37°C with agitation in a thermomixer (Eppendorf, 5382000031). The reaction was terminated by the addition of 200 µL 0.2% trifluoroacetic acid (TFA).

#### Peptide Cleanup via Solid Phase Extraction (SPE)

Following trypsin digestion, peptide mixtures underwent purification via solid-phase extraction (SPE) to attain high purity and concentration, which is essential for subsequent mass spectrometric analysis. The SPE cleanup procedure commenced with the preconditioning of Biotage SPE cartridges, which were pre-wetted twice with 1 mL of 60% acetonitrile (ACN) in 0.1% trifluoroacetic acid (TFA), ensuring adequate preparation for peptide binding. This was followed by an equilibration step in which cartridges were treated twice with 1 mL of 0.1% TFA to optimise the binding environment for the peptides. Acidified peptide samples were then loaded onto equilibrated cartridges and allowed to elute under gravity, a process designed to facilitate effective peptide binding. To further purify the captured peptides, the cartridges were washed twice with 0.1% TFA to eliminate residual impurities. Peptides were subsequently eluted into new 1.5 mL microtubes using two sequential aliquots of 500 µL of 60% ACN in 0.1% TFA, ensuring peptide recovery. The eluted peptides were then desiccated using a rotational evaporator under vacuum at room temperature for 7.5 hours, resulting in dry peptide samples that were stored at − 20° C until analysis.

#### Pre-analysis Sample Preparation and Chromatographic Conditions

The dried peptide digests were reconstituted in 100 µL of 5% ACN and 0.1% TFA. The peptide concentration was determined using the Pierce Colorimetric Peptide assay. Each sample was diluted to a final concentration of 100 ng/µL, and a pooled QC sample comprising 30 µL of each digest was prepared. Chromatographic separation was achieved on a 10 cm Acquity Premier column (2.1 mm diameter, 1.7 µm particle size) (BEH C18, 130Å, Waters, 186009658), with the column maintained at 60°C. The injection volume was 5 µL, equivalent to 500 ng of peptides on the column. The initial mobile phase conditions were 95% A (0.1% formic acid in water) and 5% B (0.1% formic acid in acetonitrile), maintained for 1 min before applying a gradient over 13.4 minutes to reach 35% B, followed by a ramp to 100% B for 1 min, and sustained for an additional 2 min. The system was subsequently returned to the initial conditions for column equilibration with a total run time of 20 min per injection.

#### Instrument Parameters

The instrument was operated in positive ion mode with a capillary voltage of 2.8 kV and a cone voltage maintained at 35 V. The source and desolvation temperatures were set at 150°C and 500°C, respectively, with desolvation and cone gas flows of 700 L/h and 150 L/h, respectively.

#### Final Method and Analysis

Samples were analysed utilising the final method, with pooled samples injected subsequent to every third sample to monitor instrument performance. Data integration was conducted using Skyline, and the results were exported to a CSV file for further analysis in Excel.

length tau in a neuronal cell model. Proc Natl Acad Sci U S A 104: 10252-10257 Doi https://doi.org/10.1073/pnas.0703676104
